# Genome reduction in an abundant and ubiquitous soil bacterial lineage

**DOI:** 10.1101/053942

**Authors:** Tess E Brewer, Kim M Handley, Paul Carini, Jack A Gibert, Noah Fierer

## Abstract

Although bacteria within the *Verrucomicrobia* phylum are pervasive in soils around the world, they are underrepresented in both isolate collections and genomic databases. Here we describe a single verrucomicrobial phylotype within the class *Spartobacteria* that is not closely related to any previously described taxa. We examined >1000 soils and found this spartobacterial phylotype to be ubiquitous and consistently one of the most abundant soil bacterial phylotypes, particularly in grasslands, where it was typically the most abundant phylotype. We reconstructed a nearly complete genome of this phylotype from a soil metagenome for which we propose the provisional name ‘*Candidatus* Udaeobacter copiosus’. The *Ca*. U. copiosus genome is unusually small for soil bacteria, estimated to be only 2.81 Mbp compared to the predicted effective mean genome size of 4.74 Mbp for soil bacteria. Metabolic reconstruction suggests that *Ca*. U. copiosus is an aerobic chemoorganoheterotroph with numerous amino acid and vitamin auxotrophies. The large population size, relatively small genome and multiple putative auxotrophies characteristic of *Ca*. U. copiosus suggests that it may be undergoing streamlining selection to minimize cellular architecture, a phenomenon previously thought to be restricted to aquatic bacteria. Although many soil bacteria need relatively large, complex genomes to be successful in soil, *Ca*. U. copiosus appears to have identified an alternate strategy, sacrificing metabolic versatility for efficiency to become dominant in the soil environment.

## Introduction

Soils harbor massive amounts of undescribed microbial diversity. For example, more than 120,000 unique bacterial and archaeal taxa were found in surface soils of Central Park in New York City, of which only ~15% had 16S rRNA gene sequences matching those contained in reference databases and <1% had representative genome sequence information (1). This undescribed soil microbial diversity is not evenly distributed across the tree of life. For example, *Acidobacteria* and *Verrucomicrobia*, two of the more abundant bacterial phyla found in soil (2, 3), represent only 0.08% and 0.06% of all cultured bacterial isolates in the Ribosomal Database Project (RDP, 4) and only 0.08% and 0.14% of publicly-available bacterial genomes found in Integrated Microbial Genomes (IMG, 5), respectively. Although the ecology and genomic attributes of abundant soil taxa are beginning to be described (6), we still lack basic information on the vast majority of soil microbes. These knowledge gaps highlight that a huge fraction of living biomass in terrestrial systems remains undescribed (7) and that we are only beginning to identify the influence of specific microbes on soil biogeochemistry and fertility.

For this study, we focus our exploration of undescribed microbial diversity on the *Verrucomicrobia* phylum. Although *Verrucomicrobia are* generally recognized as being among the most numerically abundant taxa in soil (2, 3), we know very little about the ecological or genomic attributes that contribute to their success. The phylum *Verrucomicrobia* is highly diverse and its members possess a broad range of metabolic capabilities. For example, members of the class *Methylacidiphilae* are nitrogen-fixing acidophiles capable of methane oxidation (8) while *Akkermansia muciniphila* of the class *Verrucomicrobiae* is a mucin-degrading resident of the human gut linked to reduced host obesity (9). However, the dominant *Verrucomicrobia* found in soil typically belong to the class *Spartobacteria*. For example, while *Verrucomicrobia* accounted for >50% of all bacterial 16S rRNA gene sequences in tallgrass prairie soils in the United States, >75% of these sequences were assigned to the class *Spartobacteria* (10). Currently, the class *Spartobacteria* contains only a single described and sequenced isolate, *Chthoniobacter flavus*, a slow-growing aerobic chemoorganoheterotroph capable of using common components of plant biomass for growth (11, 12). While *Spartobacteria* are prevalent in soils, they have also been observed in marine systems (‘*Spartobacteria baltica*’, 13) and as nematode symbionts (genus *Xiphinematobacter*, 14).

Here we report the distribution of a dominant *Spartobacteria* lineage, compiling data from both amplicon and shotgun metagenomic 16S rRNA gene surveys to quantify its relative abundance across >1000 unique soils. We assembled a near-complete genome of this lineage from a single soil where it was exceptionally abundant. These results provide our first glimpse into the phylogeny, ecology, and potential physiological traits of a dominant soil Verrucomicrobia and suggest that members of this group are efficient at growing and persisting in the low resource conditions common in many soil microenvironments.

## Results and Discussion

### Distribution of the dominant Verrucomicrobia in soil

A single spartobacterial clade dominates bacterial communities found in a wide range of soil types across the globe. One phylotype from this group of *Spartobacteria* represented up to 31% of total 16S rRNA gene sequences recovered from prairie soils (10). This phylotype shares 99% 16S rRNA gene sequence identity with a ribosomal clone named ‘Da101’, first described in 1998 as a particularly abundant 16S rRNA sequence recovered from a grassland soil in the Netherlands (15). To determine if the Da101 phylotype (termed ‘Da101’ herein) is abundant in other soils, we re-analyzed amplicon 16S rRNA gene sequence data obtained from >1000 soils representing a wide range of soil and site characteristics (Table S1). We found that Da101 was on average ranked within the top two most abundant bacterial phylotypes in each study (Fig. 1). In over 70% of the soils analyzed Da101 was within the top ten most abundant phylotypes. Interestingly, phylotypes belonging to the same family as Da101 (Chthoniobacteraceae) were also found within the top 5 most abundant phylotypes of several studies (Fig. 1).

**Fig. 1.**
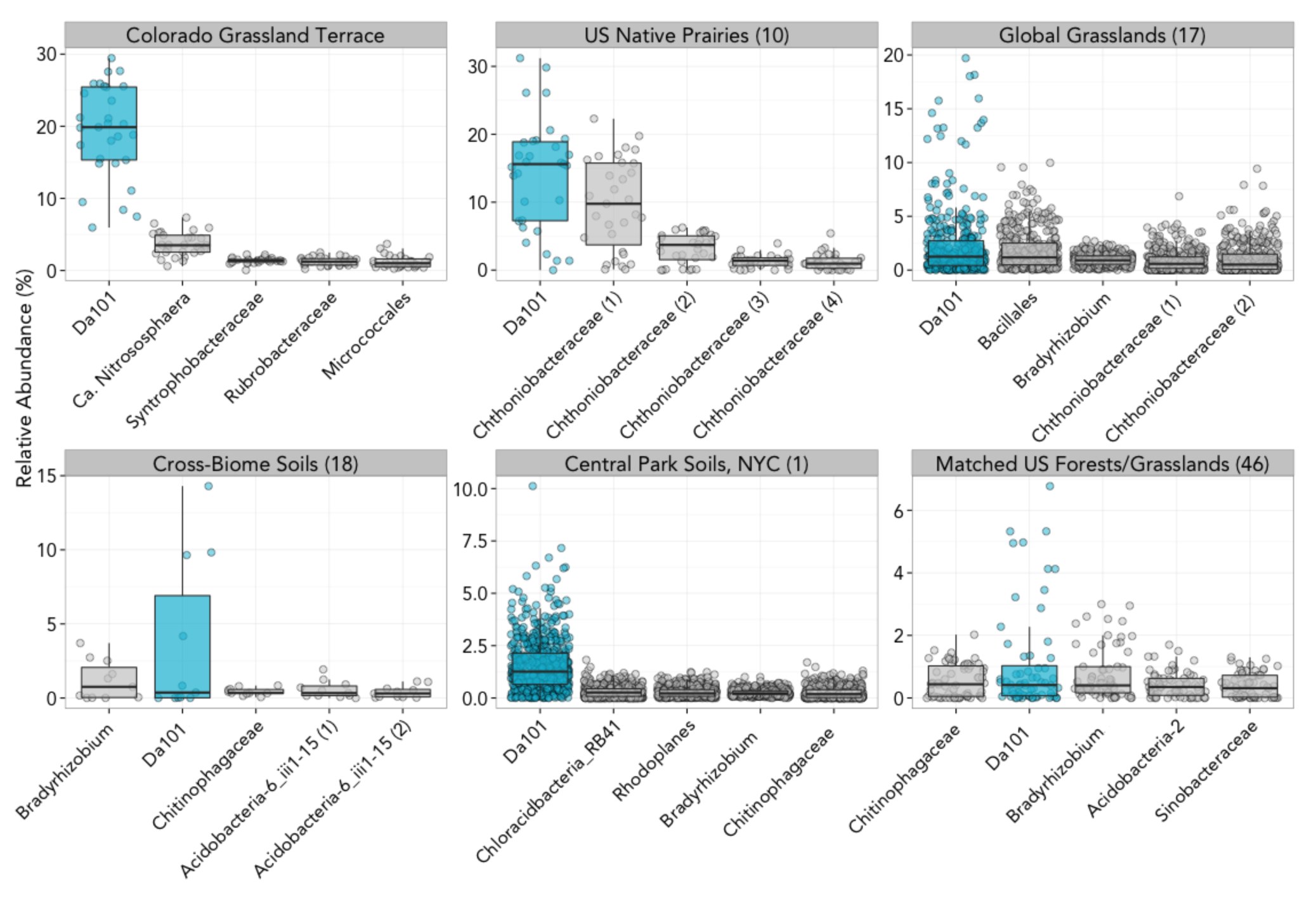
Da101 is, on average, one of the most abundant bacterial phylotypes found across >1000 soils collected from a wide range of soil and ecosystem types across the globe. The Da101 phylotype is indicated in blue while other abundant taxa are indicated in grey. Taxa are listed on the x-axis in order of their rank abundance (taxa on the left are the most abundant). Data comes from (10) Fierwer et al. 2013, (17) Leff et al. 2015, (18) Fierer et al. 2012, (1) Ramirez et al. 2014, and (46) Crowther et al. 2014. Further details on each of these studies are provided in Table S1.

As some 16S rRNA gene PCR primer sets can misestimate the relative abundance of *Verrucomicrobia* (3, 16), we investigated whether the apparent numerical dominance of Da101 in the amplicon datasets was a product of PCR primer biases. To do so, we quantified the abundance of Da101 16S rRNA genes within 75 previously published soil shotgun metagenomes (17, 18). The relative abundance of Da101 in amplicon data was reasonably well correlated with the relative abundance of Da101 determined from shotgun metagenomic data (*P*<0.0001, ρ=0.50). Confirming the amplicon-based results (Fig. 1), we found that Da101 was among the most abundant phylotypes observed in the soil bacterial communities characterized via shotgun metagenomic sequencing (Fig. S1). Thus, we conclude that the numerical dominance of Da101 in soils is not simply a product of primer biases.

Despite Da101 being one of the most abundant phylotypes found in soil, its proportional abundance can vary significantly across soil types (Fig. 1 and S1). We used metadata associated with each soil sample to determine which of the measured soil and site characteristics best predicted the relative abundance of Da101. We found that Da101 was significantly more abundant in grassland soils than in forest soils (*P*<0.0001, two tailed *t* test, Fig. S2); on average, Da101 is six times more abundant in grassland soils. These findings suggest that the soils in which Da101 excels do not overlap with those forest soils dominated by non-symbiotic *Bradyrhizobium* taxa (6). Across the grassland soils included in our meta-analysis, the relative abundance of Da101 was positively correlated with both soil microbial biomass (*P*<0.0001, ρ=0.57, Fig. S3), and aboveground plant biomass (*P*<0.0001, ρ=0.47, Fig. S3). Together, these results indicate that Da101 prefers soils receiving elevated amounts of labile carbon inputs. We did not identify any consistent correlations between the abundance of Da101 and other prokaryotic or eukaryotic taxa, suggesting that Da101 is unlikely to be a part of an obligate pathogenic or symbiotic relationship.

### Diversity of soil Verrucomicrobia

We determined the phylogenetic placement of Da101 and other soil *Verrucomicrobia* by assembling near full-length 16S rRNA gene sequences from six distinct grassland soils collected from multiple continents (Fig. 2, Table S2) using EMIRGE (19). Although we were able to assemble representative 16S rRNA gene sequences from all verrucomicrobial classes except *Methylacidiphilae*, 93% of verrucomicrobial sequences fell within the *Spartobacteria*class and 87% of these fell within the Da101 clade. These phylogenetic analyses confirm that Da101 belongs to the *Spartobacteria* class (Fig. 2). However, within the Spartobacteria class, the Da101 clade is clearly distinct from the clade containing *Chthoniobacter flavus* (11, 12), as Da101 shares only 92%16S rRNA gene sequence identity with *C. flavus*. These findings indicate that Da101 is likely a representative of a new verrucomicrobial genus. We propose the candidate genus name ‘*Candidatus* Udaeobacter’ for the Da101 clade; the proposed name combines Udaeus (‘of the earth’, Greek) with bacter (rod or staff, Greek), and like *Chthoniobacter* refers to one of the Spartoi of the Cadmus myth. We recommend the provisional name ‘*Candidatus* Udaeobacter copiosus’ for the Da101 phylotype, which refers to its numerical dominance in soil.

**Fig. 2.**
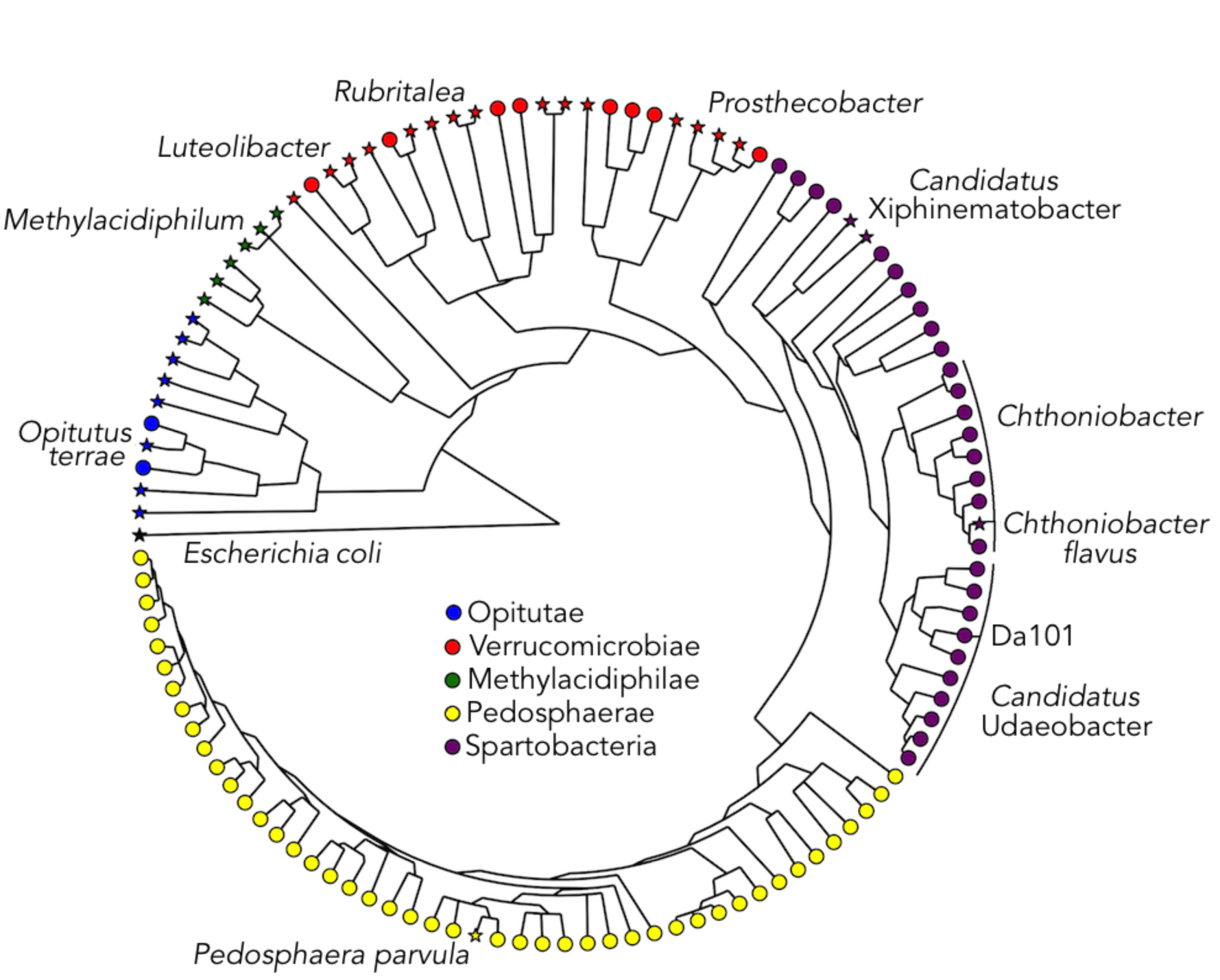
Phylogenetic analyses of soil Verrucomicrobia. Stars denote 16S rRNA gene sequences of named isolates while circles represent environmental 16S rRNA gene sequences assembled from 6 soils using EMIRGE (Table S2). The uncultivated verrucomicrobial phylotype Da101 falls within a cluster distinct from cultivated *Spartobacteria*. The UPGMA phylogenetic tree was constructed using 1200bp verrucomicrobial 16S rRNA gene sequences and is rooted with a 16S rRNA gene sequence from *Escherichia coli* K-12. Notable verrucomicrobial isolates and genera are labeled. Colors indicate verrucomicrobial classes.

### Draft genome of ‘Candidatus Udaeobacter copiosus’ recovered from metagenomic data

Despite their ubiquity and abundance in soil, there is no genomic data currently available for any representative of the ‘*Candidatus* Udaeobacter’ clade. Typically, soil hyper-diversity confounds the assembly of genomes from metagenomes (20), requiring single-cell analysis or laboratory isolation to produce an assembled genome. However, we leveraged the sheer abundance of *Ca*. U. copiosus in an individual soil to obtain a nearly complete genome from shotgun metagenomic data. We deeply sequenced a soil where *Ca*. U. copiosus accounted for >30% of 16S rRNA gene sequences and assembled a draft genome from the resulting metagenome. We used GC content, coverage, tetranucleotide frequencies, and the phylogenetic affiliation of predicted proteins to bin assembled contigs, resulting in a draft *Ca*. U. copiosus genome with 238 contigs. The draft genome is 2.65 Mbp in size, has a GC content of 54%, and encodes for 3,042 predicted proteins, 67% of which could be assigned to Pfam protein families (21) by the IMG annotation pipeline (5). We estimate that the full *Ca*. U copiosus genome is 2.81 Mbp in length based on the recovery of 94% of single copy housekeeping genes (34 of 36 genes, Table S3) that are commonly used to estimate genome completion (22). The *Ca*. U. copiosus genome shares only 69.3% average nucleotide identity (23) with the genome of its closest sequenced relative *C. flavus*, further supporting its proposed placement as the distinct genus ‘*Candidatus* Udaeobacter’. Additionally, the *Ca*. U copiosus 16S rRNA gene has 100% identity to the Da101 phylotype sequence, indicating that this genome is indeed a representative of the aforementioned dominant Da101 clade.

*Ca*. U. copiosus has a particularly small genome size compared to *C. flavus* (2.81 Mbp to 8.80 Mbp, predicted genome sizes). To see how the genome of *Ca*. U. copiosus compares to other soil bacteria, we compiled data from 378 soil bacteria with finished or permanent draft genome sequences in IMG whose 16S rRNA gene sequences matched the 16S rRNA gene amplicon sequences obtained by Leff et al. (2015) with at least 99% identity. Nearly all (99%) of these 378 genomes came from cultivated taxa. We estimated the genome completeness for each of the 378 taxa using the same method as for *Ca*. U. copiosus and found the mean estimated genome size of these taxa to be 5.28 ± 2.15 Mbp (mean ± SD), which is nearly identical to metagenomic based estimates of mean genome size for soil microbes (24). Strikingly, the 2.81 Mbp genome of *Ca*. U. copiosus is ~50% smaller than the mean genome size of these 378 taxa and only 13% of these genomes were smaller than the genome of *Ca*. U. copiosus.

Although soil bacteria with larger genomes tend to be more common in soil, *Ca*. U. copiosus is a notable exception to this pattern. We linked the genome size of each of the matched IMG bacterial genomes with the average abundance of their corresponding amplicon sequence from Leff et al. (2015) and found that genome size is positively correlated with average relative abundance (*P* <0.001, ρ=0.37, Fig. 3). That is, bacteria with large genomes tend to comprise a significantly larger proportion of soil bacterial communities. On average, the genomes of soil prokaryotes are larger than those inhabiting aquatic ecosystems (25) or the human gut (26). These relatively large genomes are thought to provide soil-dwelling bacteria with a more diverse genetic inventory to enhance survival in conditions where resources are diverse, but sparse (27, 28). However, the *Ca*. U. copiosus genome has a conspicuously reduced genome given its numerical abundance (Fig. 3). This suggests that *Ca*. U. copiosus occupies a niche space that does not require expansive functional diversity and points to an alternative route to success for soil bacteria. These results also suggest that abundant, uncultivated soil bacteria may have smaller genomes than the cultivated taxa that represent the vast majority of available genomic data. A similar pattern has been observed in aquatic systems where uncultivated taxa often have smaller genomes than cultivated taxa (29). Because the majority of available genomic information is derived from cultivated bacterial taxa, the lack of genomic information from bacteria with reduced genomes likely stems from challenges associated with culturing taxa with reduced genomes (25).

**Fig. 3.**
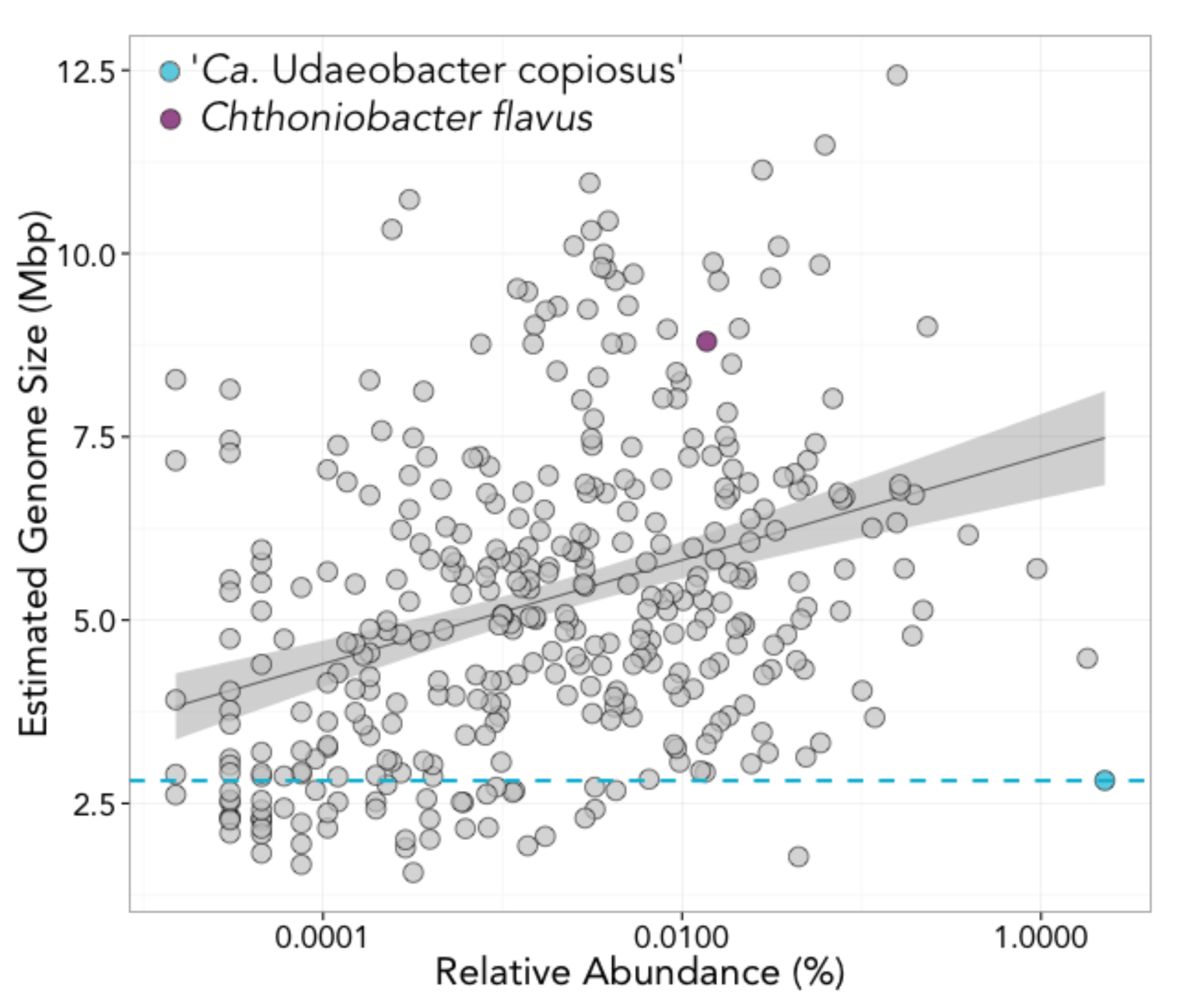
‘*Ca*. Udaeobacter copiosus’ has a reduced genome size compared to other abundant soil bacteria. Points represent the estimated genome size and relative abundances of 378 bacterial genomes obtained by matching 16S rRNA gene sequences from (17) to 16S rRNA gene sequences extracted from genomes in IMG at 99% sequence identity. Only genomes classified as ‘permanent draft’ or ‘finished’ status were used. Bacteria with larger genomes tend to be more abundant (p<0.0001, ρ=0.368, Spearman correlation) with *Ca*. U. copiosus (indicated in blue) being a notable exception to this pattern, as it has a high relative abundance (2.26% of 16S rRNA sequences) but has a relatively small genome.

Metabolic reconstruction of the *Ca*. U. copiosus genome points to an aerobic chemoorganoheterotrophic lifestyle with the capacity to use a limited range of carbon substrates for growth including glucose, pyruvate, and chitobiose. Glycogen/starch synthesis and utilization genes were identified (*glgABCP* and *amyA*), suggesting that *Ca*. U. copiosus has the capacity to store surplus carbon as glycogen or starch. Glycogen metabolism has been demonstrated in other *Verrucomicrobia* (30). Genes encoding for the complete biosynthesis of vitamins B_2_, B_3_, B_5_ (from valine) and B_6_ were recovered. Full biosynthetic pathways for *de novo* synthesis of alanine, aspartate, asparginine, glutamate, glutamine and proline were also present. Nearly complete pathways were recovered for glycine, threonine and methionine biosynthesis. Genes encoding for the conversion of methionine to cysteine were present as the only apparent route to cysteine biosynthesis. The only putative serine biosynthesis pathway is via the transamination of pyruvate. Genes indicative of autotrophic metabolism (for example, RuBisCO, ATP citrate lyase) were not identified. Additionally, genes indicative of methanotrophy (*pmo*), methylotrophy (*mxaF* or *xoxF*), ammonia (*amo*) or nitrite oxidation (*nxr*) were absent.

Genes encoding for the biosynthesis of all branched-chain (isoleucine, leucine and valine) and aromatic (tryptophan, tyrosine and phenylalanine) amino acids were conspicuously absent from the *Ca*. U copiosus genome. Additionally, the biosynthetic pathways for arginine and histidine were also absent. These eight amino acids are among the most energetically expensive to make (Fig S4, 31), suggesting that their acquisition from the environment offers *Ca*. U copiosus an energetic savings relative to taxa that synthesize them *de novo*. In contrast, Ca. U. copiosus does have the complete suite of genes for the biosynthesis of those amino acids that are energetically less expensive to make (including alanine, aspartate, and glutamate, Fig. S4). The absence of branched-chain amino acid synthesis pathways in the *Ca*. U. copiosus genome is consistent with previous observations that these genes are underrepresented in natural populations of soil *Verrucomicrobia* (10). Although *Ca*. U. copiosus lacks many amino acid synthesis pathways, numerous genes encoding for peptide transport, degradation and recycling were identified. For example, when scaled for genome size, *Ca*. U. copiosus encodes four times as many putative peptide and amino acid transporters as *C. flavus* (1.5% of genome to 0.37%) and twice as many predicted proteases (6.5% of genome versus 3.2%). *Ca*. U. copiosus also encodes for all components of the bacterial proteasome. Proteasomal degradation is critical for amino acid recycling under starvation conditions in mycobacteria (32). The enrichment of peptide transport and degradation systems in the *Ca*. U. copiosus genome suggest that at least some of the amino acids *Ca*. U. copiosus cannot synthesize are available directly from the soil environment or by indirect associations with other soil biota. In addition to these likely amino acid auxotrophies, *Ca*. U. copiosus has several putative B-vitamin or B-vitamin precursor requirements. For example, the entire vitamin B_12_ synthesis pathway was absent in *Ca*. U. copiosus, despite the presence of three genes encoding vitamin B_12_-dependent proteins (methionine synthase, ribonucleotide reductase, and methylmalonyl-CoA mutase). Vitamin B_12_ auxotrophies are relatively common in soil (33), making it likely that exogenous vitamin B_12_ is generally available to many soil bacteria. Genes encoding for the complete vitamin B_1_ biosynthetic pathway was complete except for the 4-amino-3-hydroxymethyl-2 methylpyrmidine (HMP) synthase (*thiC*), encoding the first step in B_1_ biosynthesis. The absence of this gene, in the presence of the remainder of the B_1_ synthesis pathway, was recently linked to an obligate HMP requirement in marine bacteria (34).

The abundance and cosmopolitan distribution of *Ca*. U. copiosus (Fig. 1), together with its small genome size relative to other soil microbes (Fig. 3), suggest that it is undergoing streamlining selection to minimize genome size. The genome-streamlining hypothesis proposes that, in large bacterial populations, reduced genome complexity is a trait under natural selection, especially in environments where nutrients are sparse and can periodically limit growth (25). All contemporary free-living organisms with streamlined genomes inhabit aquatic environments (25, 35). However, compared to these aquatic environments, soils are more heterogeneous (36), have higher overall microbial diversity (37), and slower carbon turnover (38). Therefore, the functional complexity required by soil microbes to succeed within a given niche is likely large relative to that required by aquatic microbes. This means that the effects of genome streamlining are likely to be most evident (i.e., result in smaller genomes) in aquatic environments and that we might expect genome reduction to be relatively uncommon across soil taxa. This expectation is consistent with the fact that, on average, the genomes of aquatic microbes are smaller than their terrestrial counterparts (25). However, the small genome and numerous putative auxotrophies of *Ca*. U. copiosus show that genome streamlining is not unique to aquatic organisms and that genome streamlining may also confer a selective growth advantage in the soil environment.

Genome streamlining in *Ca*. U. copiosus has resulted in reduced catabolic and biosynthetic capacity, and thus a loss of metabolic versatility. The absence of multiple costly amino acid and vitamin biosynthetic pathways from the *Ca*. U. copiosus genome implies that these compounds can be acquired from the soil environment. Several studies have shown that free amino acids are present in soil (39, 40), although oligopeptides are reported to be more abundant in grasslands and may be assimilated with kinetics similar to free amino acids (41). The enrichment of proteases and amino acid and peptide importers in the *Ca*. U. copiosus genome suggests that it is well equipped to assimilate this fraction of soil organic matter. Dispensing the capacity to synthesize costly amino acids and vitamins likely provides *Ca*. U copiosus a growth advantage in resource limiting conditions when competition for labile carbon is high. Alternatively, many of the putative amino acid auxotrophies described here are involved in synergistic growth (42) and may be supplied by other microbes as common community goods (43). Based on the few spartobacterial isolates that have been cultivated (11), culture-independent studies (10, 44), and the genomic data presented here, we speculate that *Ca*. U. copiosus is a small, oligotrophic soil bacterium that reduces its requirement for soil organic carbon by acquiring costly amino acids and vitamins from the environment.

### Conclusions

Whereas successful soil microbes are predicted to have large genomes (27, 28, Fig. 3), *Ca*. U. copiosus has a small genome, indicating that, similar to aquatic microbes, minimization of cellular architecture can also represent a successful strategy for soil microbes. We do not know if other uncultivated abundant soil taxa also contain streamlined genomes because pre-existing genome databases are preferentially biased towards cultivated isolates. For example, only 4.5% of bacterial genomes in IMG are from uncultivated taxa (accessed April 2016). Bacteria encoding for greater metabolic versatility likely have larger genomes and therefore may be easier to culture in the laboratory (29). On the other hand, specific and combinatorial nutrient requirements such as those described for *Ca*. U. copiosus, present a complex problem for researchers attempting to cultivate microbes with reduced genomes (45). Although *Ca*. U. copiosus has not yet been grown in the laboratory, cultivation is clearly a crucial next step to describing this organism, using the information described here to ‘tailor’ a growth medium specifically for *Ca*. U. copiosus and related microbes. Such an approach will undoubtedly improve our ability to describe and study the majority of soil microbes, even dominant soil microbes like *Ca*. U. copiosus, which remain difficult to cultivate under laboratory conditions.

## Materials and Methods

### Estimating the abundances and distributions of Verrucomicrobia in soil

While five abundant *Verrucomicrobia* phylotypes were described in Fierer et al. (2013), a single phylotype with 99% identity to the clone Da101 (15) was clearly dominant. We searched previously published soil datasets for representative sequences with 100% identity to this Da101 phylotype, including 31 soils from United States native tallgrass prairies (10), 64 soils from matched forest and grassland sites across North America (46), 595 soils collected from Central Park in New York City (1), 367 grassland soils collected from North America, Europe, Australia, and Africa (17), and a cross-biome collection of 11 desert and non-desert soils from across the globe (18). We also included a dataset from a grassland terrace near Boulder, Colorado (105.23W, 40.12N, Table Mountain) where 30 soils were collected from a depth of 25 cm within a 100m^2^ area on 28Jan15. Amplicon sequences and associated metadata from this study are available at https://dx.doi.org/10.6084/m9.figshare.3363505.v3. Collectively these datasets represent 1097 unique soil samples collected from a wide range of ecosystem and soil types.

For all samples, DNA was extracted with the MoBio PowerSoil kit and the V4 region of the 16S rRNA gene was amplified in triplicate with the 515f/806r primer pair. After normalization to equimolar concentrations, amplicons were sequenced on an Illumina MiSeq (151bp paired end) at the University of Colorado BioFrontiers Institute Next-Gen Sequencing Core Facility. Sequences were processed as described previously (17). In brief, we used a combination of QIIME (47) and UPARSE (48) to quality-filter, remove singletons, and merge paired reads. Sequences were assembled into phylotypes at the 97% identity level using UCLUST (49). Taxonomy was assigned using the Greengenes 13_8 database (50) and the Ribosomal Database Project classifier (4) and each dataset was rarefied independently (Table S1).

As PCR primer biases can misestimate the relative abundances of *Verrucomicrobia* (3,16), we also estimated the abundances of the Da101 phylotype directly from shotgun metagenomic data. We used Metaxa2 with default settings (51) to extract bacterial 16S rRNA gene sequences from shotgun metagenomic data compiled from previous analyses of 75 different soils (data from 17, 18). Extracted 16S rRNA gene fragments were matched to GreenGenes full-length sequences at 99% ID using the usearch7 command usearch_global. The matched Greengenes sequences were then clustered and assigned taxonomy as described above.

### Describing the phylogenetic diversity of soil Verrucomicrobia

We reconstructed near-full length 16S rRNA gene sequences to construct a phylogeny of soil *Verrucomicrobia* from six soil samples (see Table S2) that were selected to represent geographically distinct grasslands with a range of verrucomicrobial abundances. We extracted DNA from each of these soils as described previously (17) and used the 27f/1392r primer pair (52) to amplify near full-length 16S rRNA genes as described in (19). The amplicons were sheared using the Covaris M220 (Covaris, Woburn, MA) and the 16S rRNA gene libraries were prepared using TruSeq DNA LT library preparation kits (Illumina, San Diego, CA). Samples were pooled and sequenced on an Illumina MiSeq (2x300bp) at the University of Colorado Next Generation Sequencing Facility.

After quality filtering of sequences, near full length SSU sequences were reconstructed using EMIRGE (19). After 40 iterations, sequences were merged into phylotypes with ≥97% similarity. Reconstructed sequences were trimmed to 1200 bp and all sequences were further clustered at 95% identity due to gaps in some assemblies. Full-length 16S rRNA sequences from named verrucomicrobial isolates were aligned along with the reconstructed sequences using PyNAST (53). A UPGMA tree was constructed using the R packages seqnir and phangorn and visualized with GraPhlAn (R 3.2.2, version 0.9.7).

### Assembly and annotation of a genome from the dominant soil Verrucomicrobial phylotype

We assembled the genome of ‘*Candidatus* Udaeobacter copiosus’ from a metagenome of a U.S. prairie soil sample (NTP21, Hayden, IA) estimated to have particularly high abundances of bacteria within the Da101 clade (10). Fragmented DNA extracted from this soil was prepared for sequencing using WaferGen’s PrepX ILM DNA library Kit (WaferGen Biosystems Inc, Fremont, CA) and the Apollo 324 Automated Library Prep System for library generation. The library was sequenced on one Illumina HiSeq2000 lane (2×101 bp), yielding 17 Gb of sequence with an average paired-end insert size of 345 bp. Low quality reads were trimmed using Sickle v. 1.29 with a quality score threshold of Q=3, or removed if trimmed to <80 bp long (https://github.com/najoshi/sickle). The sequences were assembled using IDBA_ud v. 1.1.0 (54) with a kmer range of 40 to 70 and step size of 15. To improve recovery of the most abundant *Verrucomicrobia*, the genome was selectively re-assembled using Velvet with a kmer size of 59, and expected kmer coverage of 11.5 (range 7.5 to 15.5). To bin contigs ≥2 kb long, genes and protein sequences were predicted using Prodigal v. 2.60 in metagenomics mode (55). For each contig, we determined the GC content, coverage, and the phylogenetic affiliation based on the best hit for each predicted protein in the Uniref90 database (Sept-2013, 56) following ublast searches (49). We also constructed emergent self-organizing maps (ESOM) using tetranucleotide frequencies of 5 kb DNA fragments (57). A combination of these approaches were used to identify the genome. The draft genome was uploaded to IMG for annotation under the taxon ID 2651869889.

No rRNA genes were annotated by IMG, so we used Metaxa with default settings on the unassembled sequences to extract any 16S rRNA genes. Metaxa recovered two ~500 bp 16S rRNA gene fragments at 23-29× coverage which aligned to separate regions of the full-length 16S rRNA gene from the closest related verrucomicrobial genome (*C. flavus*). Because these two rRNA gene fragments have the same coverage as the genome and align to separate regions of one 16S rRNA gene, it is likely that the sequences encode a single rRNA operon.

## Acknowledgements

Funding to support this work was provided by grants from the National Science Foundation to N.F. (DEB0953331, EAR1331828), and a Visiting Postdoctoral Fellowship award to P.C. from the Cooperative Institute for Research in Environmental Sciences.

---

**Fig. S1:**
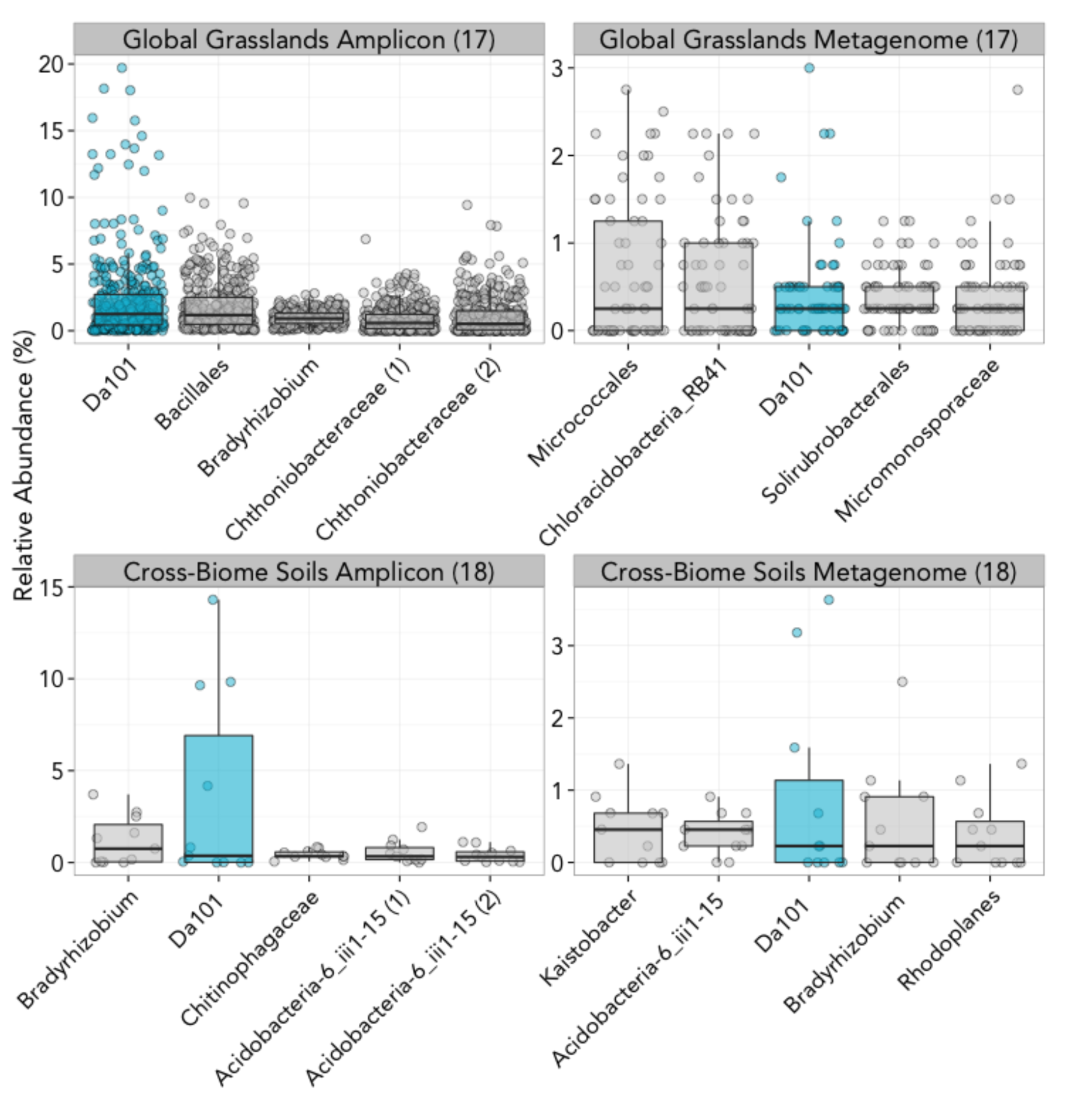
Da101 rank is similar in amplicon and metagenomic data. The top 5 phylotypes from two matched amplicon and metagenomic datasets (Leff et al. 2015 (17), Fierer et al. 2012 (18)) are shown in order of decreasing median rank. Each point represents one sample within the corresponding dataset. Da101’s position is highlighted with blue while all other phylotypes are grey.

**Fig. S2.**
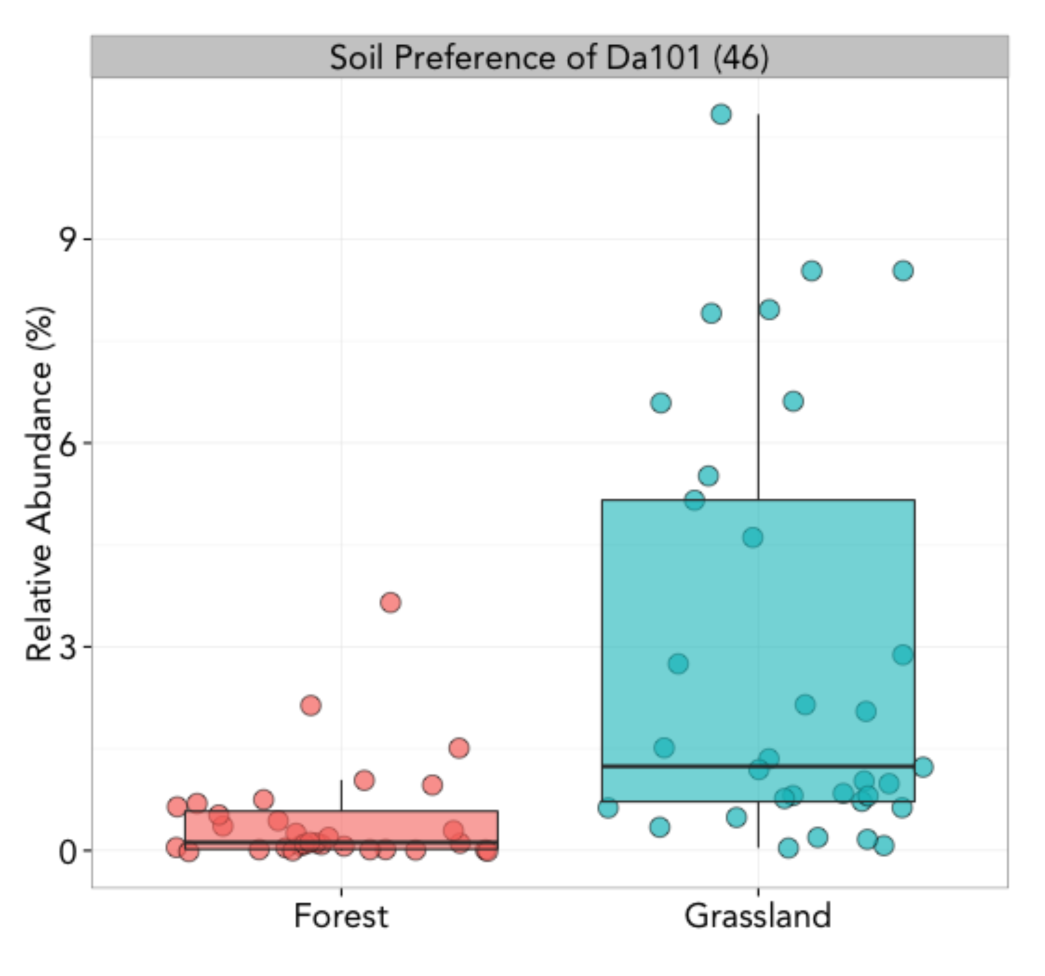
Phylotype Da101 is more abundant in grasslands than forests; p<0.0001, two tailed t-test. Data is from Crowther et al. 2014 (46).

**Fig. S3.**
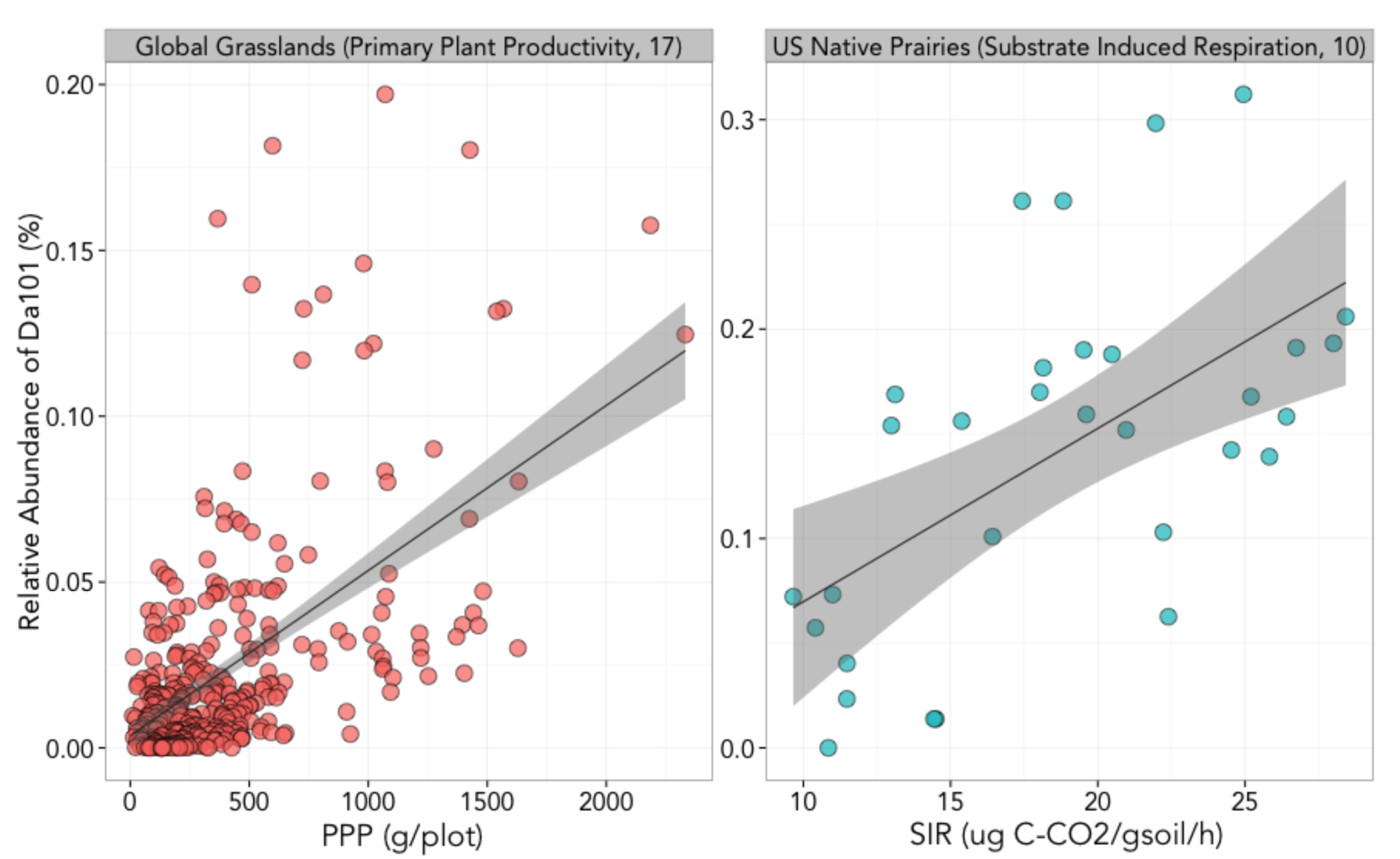
The abundance of phylotype Da101 correlates with measures of microbial and plant biomass in two separate studies: SIR p<0.001 ρ=0.57, PPP p<0.0001 ρ=0.47, Spearman correlations. Data is from Fierer et al. 2013 (10) and Leff et al. 2015 (17).

**Fig. S4.**
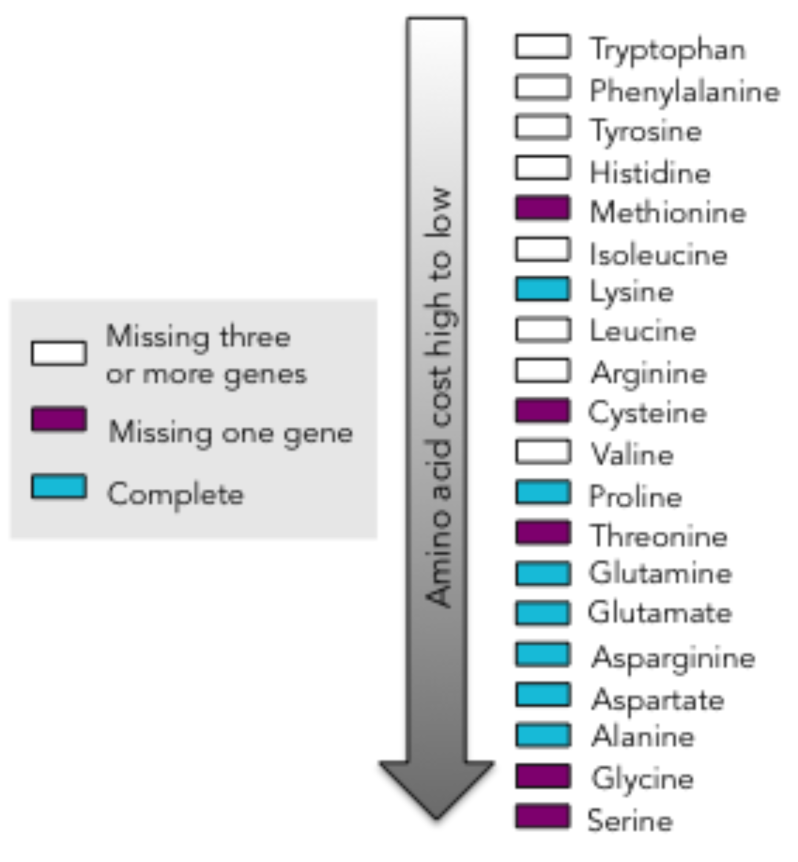
‘*Ca*. Udaeobacter copiosus’ lacks pathways to synthesize several expensive amino acids, yet has partial or complete pathways for relatively inexpensive amino acids. Cost of each of amino acid was estimated in *E. coli* by number of high-energy phosphate bonds hydrolyzed (31).

**Table S1:** Descriptions of Datasets Used in this Study

|  | Fierer<br>et al. 2012 | Fierer<br>et al. 2013 | Crowther<br>et al. 2014 | Ramirez<br>et al. 2014 | Leff<br>et al. 2015 | Table<br>Mountain, CO |
| --- | --- | --- | --- | --- | --- | --- |
| **REGION DESCRIPTION** | Global<br>cross-biome | U.S. native<br>prairies | U.S. matched<br>forest/grasslands | Central<br>park, NYC | Global<br>grasslands | CO<br>Terrace |
| **# SAMPLES** | 11 | 31 | 64 | 595 | 367 | 29 |
| **# SITES** | 11 | 31 | 11 | 1 | 25 | 1 |
| **METAGENOMIC DATA** | 11 | - | - | - | 87 | - |
| **ITS DATA** | × | × | ✓ | ✓ | ✓ | × |
| **18S DATA** | × | × | × | ✓ | × | × |
| **pH RANGE** | 4.1-9.5 | 5.8-7.9 | 4.0-8.1 | 3.9-8.4 | 4.4-8.2 | - |
| **MAT RANGE (°C)** | -19-25 | 3.7-18.9 | -3.2-22.8 | 13 | 0.3-18.4 | 11 |
| **MAP RANGE (mm)** | 100-4000 | 503-1148 | 287-3460 | 1016 | 262-1898 | 525 |
| **RAREFACTION DEPTH** | 15000 | 942 | 4000 | 5000 | 18000 | 11000 |

**Table S2:** Samples used to construct EMIRGE phylogeny

| **SAMPLE** | **DATASET** | **LOCATION** | **LATITUDE** | **LONGITUDE** | **DESCRIPTION** |
| --- | --- | --- | --- | --- | --- |
| NTP21 | Fierer et al. 2013 | Hayden, IA | 43.26 | -92.23 | Native prairie |
| NTP28 | Fierer et al. 2013 | Glynn Prairie, MN | 44.15 | -95.41 | Native prairie |
| NN1182 | Leff et al. 2015 | Val Mustair, Switzerland | 46.63 | 10.37 | Alpine grassland |
| NN772 | Leff et al. 2015 | Msunduzi Municipality, South Africa | -29.67 | 30.40 | Mesic grassland |
| TM25 | New data set | Table Mountain, CO | 40.01 | -105.5 | Grassland terrace |
| GG14 | New data set | Gordon Gulch, CO | 40.12 | -105.2 | Meadow |

**Table S3:** ‘*Candidates* Udaeobacter copiosus’ genome has 34/36 single copy housekeeping genes

| IMG GENE ID | COG/PFAM PRODUCT NAME | SEQUENCE LENGTH (BP) | COG/PFAM |
| --- | --- | --- | --- |
| 2653240560 | arginyl-tRNA synthetase | 1737 | COG0018 |
| 2653239845 | DNA-directed RNA polymerase subunit alpha | 1044 | COG0202 |
| 2653239819 | DNA-directed RNA polymerase subunit beta | 2910 | COG0085 |
| 2653240331 | histidyl-tRNA synthetase | 1239 | COG0124 |
| 2653239579 | Ribosome-binding ATPase GTP1/OBG family | 1125 | COG0012 |
| 2653241119 | isoleucyl-tRNA synthetase | 2748 | COG0060 |
| 2653239815 | large subunit ribosomal protein L1 | 702 | COG0081 |
| 2653239810 | large subunit ribosomal protein L11 | 429 | COG0080 |
| 2653239936 | large subunit ribosomal protein L13 | 438 | COG0102 |
| 2653241207 | large subunit ribosomal protein L14 | 366 | COG0093 |
| 2653241203 | large subunit ribosomal protein L16 | 426 | COG0197 |
| 2653241213 | large subunit ribosomal protein L18 | 375 | COG0256 |
| 2653241195 | large subunit ribosomal protein L3 | 726 | COG0087 |
| 2653241210 | large subunit ribosomal protein L5 | 573 | COG0094 |
| 2653241212 | large subunit ribosomal protein L6 | 540 | COG0097 |
| 2653239619 | leucyl-tRNA synthetase | 2412 | COG0495 |
| 2653242017 | N6-L-threonylcarbamoyladenine synthase | 1086 | COG0533 |
| 2653241883 | phenylalanyl-tRNA synthetase alpha chain | 282 | pfam01409 |
| 2653241216 | preprotein translocase subunit SecY | 1509 | COG0201 |
| 2653241215 | ribosomal protein L15 | 636 | COG0200 |
| 2653241201 | ribosomal protein L22 | 480 | COG0091 |
| 2653241222 | small subunit ribosomal protein S11 | 630 | COG0100 |
| 2653241191 | small subunit ribosomal protein S12 | 435 | COG0048 |
| 2653241221 | small subunit ribosomal protein S13 | 396 | COG0099 |
| 2653241789 | small subunit ribosomal protein S15 | 177 | pfam00312 |
| 2653241206 | small subunit ribosomal protein S17 | 297 | COG0186 |
| 2653239969 | small subunit ribosomal protein S2 | 702 | COG0052 |
| 2653241202 | small subunit ribosomal protein S3 | 690 | COG0092 |
| 2653241223 | small subunit ribosomal protein S4 | 612 | COG0522 |
| 2653241214 | small subunit ribosomal protein S5 | 564 | COG0098 |
| 2653241192 | small subunit ribosomal protein S7 | 474 | COG0049 |
| 2653241211 | small subunit ribosomal protein S8 | 399 | COG0096 |
| 2653239935 | small subunit ribosomal protein S9 | 396 | COG0103 |
| 2653241990 | valyl-tRNA synthetase | 816 | pfam00133 |

## References

1. Ramirez KS et al. (2014) Biogeographic patterns in below-ground diversity in New York City’s Central Park are similar to those observed globally. Proc R Soc B 281(1795):20141988.

2. Janssen PH. (2006) Identifying the dominant soil bacterial taxa in libraries of 16S rRNA and 16S rRNA genes. Appl Environ Microbiol 72(3):1719–28.

3. Bergmann GT et al. (2011) The under-recognized dominance of Verrucomicrobia in soil bacterial communities. Soil Biol Biochem 43(7):1450–5.

4. Wang, Q, Garrity GM, Tiedje JM, and Cole JR. (2007) Naïve Bayesian Classifier for Rapid Assignment of rRNA Sequences into the New Bacterial Taxonomy. Appl Environ Microbiol 73(16):5261–7.

5. Markowitz VM et al. (2014) IMG 4 version of the Integrated Microbial Genomes comparative analysis system. Nucleic Acids Res 42:D560–7.

6. VanInsberghe, D et al. (2015) Non-symbiotic Bradyrhizobium ecotypes dominate North American forest soils. ISME J 9(11):2435–41.

7. Fierer, N., M.S. Strickland, D. Liptzin, M.A. Bradford, C.C. Cleveland. (2009) Global patterns in belowground communities. Ecol Lett 12(11):1238–49.

8. Dunfield PF et al. (2007) Methane oxidation by an extremely acidophilic bacterium of the phylum Verrucomicrobia. Nature 450(7171):879–82.

9. Everard A et al. (2013) Cross-talk between *Akkermansia muciniphila* and intestinal epithelium controls diet-induced obesity. Proc Natl Acad Sci USA 110(22):9066–71

10. Fierer N et al. (2013) Reconstructing the microbial diversity and function of pre-agricultural tallgrass prairie soils in the United States. Science 342(6158): 621–4.

11. Sangwan P, Chen X, Hugenholtz P and Janssen PH. (2004) *Chthoniobacter flavus* gen. nov., sp. nov., the First Pure-Culture Representative of Subdivision Two, *Spartobacteria* classis nov., of the Phylum *Verrucomicrobia*. Appl Environ Microbiol 70(10):5875–81.

12. Kant R, van Passel MWJ, Palva A, et al. (2011) Genome Sequence of Chthoniobacter flavus Ellin428, an Aerobic Heterotrophic Soil Bacterium. J Bacteriol 193(11):2902–3.

13. Herlemann DPR et al. (2013) Metagenomic De Novo Assembly of an Aquatic Representative of the Verrucomicrobial Class Spartobacteria. mBio 4(3):e00569–12

14. Vandekerckhove TT, Willems A, Gillis M, Coomans A. (2000) Occurrence of novel verrucomicrobial species, endosymbiotic and associated with parthenogenesis in Xiphinema americanum-group species (Nematoda, Longidoridae). Int J Syst Evol Microbiol 50:2197–205.

15. Felske A, Akkermans ADL. (1998) Prominent occurrence of ribosomes from an uncultured bacterium of the Verrucomicrobiales cluster in grassland soils. Lett Appl Microbiol 26(3):219–23.

16. Guo J, Cole JR, Zhang Q, Brown CT, Tiedje JM. (2016) Microbial community analysis with ribosomal gene fragments from shotgun metagenomes. Appl Environ Microbiol 82(1):157–66.

17. Leff, JW et al. (2015) Consistent responses of soil microbial communities to elevated nutrient inputs in grasslands across the globe. Proc Natl Acad Sci USA 112(35):10967–72.

18. Fierer N et al. (2012) Cross-biome metagenomic analyses of soil microbial communities and their functional attributes. Proc Natl Acad Sci USA 109(52):21390–5.

19. Miller CS, Baker BJ, Thomas BC, Singer SW, Banfield JF. (2011) EMIRGE: reconstruction of full-length ribosomal genes from microbial community short read sequencing data. Genome Biol 12:R44.

20. Howe AC et al. (2014) Tackling soil diversity with the assembly of large, complex metagenomes. Proc Natl Acad Sci USA 111(13):4904–9.

21. Finn RD et al. (2016) The Pfam protein families database: towards a more sustainable future. Nucleic Acids Res 44(D1):D279–85.

22. Ciccarelli FD et al. (2006) Toward Automatic Reconstruction of a Highly Resolved Tree of Life. Science 311(5765):1283–7.

23. Varghese NJ et al. (2015) Microbial species delineation using whole genome sequences. Nucleic Acids Res 10.1093.

24. Raes J, Korbel JO, Lercher MJ, von Mering C, Bork P. (2007). Prediction of effective genome size in metagenomic samples. Genome Biol 8(1):R10.

25. Giovannoni SJ, Thrash JC, Temperton B. (2014) Implications of streamlining theory for microbial ecology. ISME J 8:1553–65.

26. Nayfach S, Pollard KS. (2015) Average genome size estimation improves comparative metagenomics and sheds light on the functional ecology of the human microbiome. Genome Biol 16(1):51.

27. Konstantinidis KT, Tiedje JM. (2003) Trends between gene content and genome size in prokaryotic species with larger genomes. Proc Natl Acad Sci USA 101(9):3160–5.

28. Barberán A et al. (2014) Why are some microbes more ubiquitous than others? Predicting the habitat breadth of soil bacteria. Ecol Lett 17(7):794–802.

29. Button DK, Robertson, BR. (2001) Determination of DNA content of aquatic bacteria by flow cytometry. Appl Environ Microbiol 67(4):1636–45.

30. Khadem AF et al. (2012) Genomic and Physiological Analysis of Carbon Storage in the Verrucomicrobial Methanotroph “*Ca*. Methylacidiphilum Fumariolicum” SolV. Front Microbiol 3:345.

31. Akashi H, and Gojobori T (2002) Metabolic efficiency and amino acid composition in the proteomes of *Escherichia coli* and *Bacillus subtilis*. Proc Natl Acad Sci USA 99(6):3695–3700.

32. Elharar Y et al. (2014) Survival of mycobacteria depends on proteasome-mediated amino acid recycling under nutrient limitation. EMBO J 33(16):1802–14.

33. Lochhead, AG. (1958) Soil bacteria and growth promoting substances. Bacteriol Rev 22(3):145–53.

34. Carini P et al.1 (2014) Discovery of a SAR11 growth requirement for thiamin’s pyrimidine precursor and its distribution in the Sargasso Sea. ISME J 8:1727–38.

35. Kantor RS et al. (2013) Small Genomes and Sparse Metabolisms of Sediment-Associated Bacteria from Four Candidate Phyla. mBio 4(5):e00708–13.

36. Vos M, Wolf AB, Jennings SJ, Kowalchuk GA. (2013) Micro-scale determinants of bacterial diversity in soil. FEMS Microbiol Rev 37(6):936–54.

37. Fierer N, Lennon JT. (2011) The generation and maintenance of diversity in microbial communities. Am J Bot 98(3):439–48.

38. Giovannoni SJ, Vergin VL. (2012) Seasonality in Ocean Microbial Communities. Science 335(6069):671–6.

39. Friedel JK, Scheller E. (2002) Composition of hydrolysable amino acids in soil organic matter and soil microbial biomass. Soil Bio Biochem 34(3):315–25.

40. Sauheitl L, Glaser B, Dippold M, Lieber K, Weigelt A (2010) Amino acid fingerprint of a grassland soil reflect changes in plant species richness. Plant Soil 334(1):353–63.

41. Farrell M et al. (2013) Oligopeptides Represent a Preferred Source of Organic N Uptake: A Global Phenomenon? Ecosystems 16(1):133–45.

42. Mee MT, Collins JJ, Church GM, Wang HH (2014) Syntrophic exchange in synthetic microbial communities. Proc Natl Acad Sci USA 111(20):E2149– E2156.

43. Morris JJ, Lenski RE, Zinser ER (2012) The Black Queen Hypothesis: Evolution of dependencies through adaptive gene loss. mBio 3(2):e00036–12.

44. Portillo, M.C., J.W. Leff, C.L. Lauber, N. Fierer. (2013) Cell size distributions of soil bacterial and archaeal taxa. Appl Environ Microbiol 79(24):7610–7617.

45. Carini P, et al. (2013) Nutrient requirments for growth of the extreme oligotroph *‘Candidatus* Pelagibacter ubique’ HTCC1062 on a defined medium. ISME J 7(3):592–602.

46. Crowther, TW et al. (2014) Predicting the responsiveness of soil biodiversity to deforestation: a cross-biome study. Glob Chang Biol 20(9):2983–94

47. Caporaso JG et al. (2010) QIIME allows analysis of high-throughput community sequencing data. Nat Methods 7(5):335–336.

48. Edgar RC (2013) UPARSE: highly accurate phylotype sequences from microbial amplicon reads. Nat Methods 10(10):996–8

49. Edgar, RC (2010) Search and clustering orders of magnitude faster than BLAST. Bioinformatics 26(19):2460–1.

50. DeSantis, T.Z et al. (2006) Greengenes, a Chimera-Checked 16S rRNA Gene Database and Workbench Compatible with ARB. Appl Environ Microbiol 72(7):5069–72.

51. Bengtsson-Palme J et al. (2015) Metaxa2: improved identification and taxonomic classification of small and large subunit rRNA in metagenomic data. Mol Ecol Resour 15(6):1403–14.

52. Miller CS et al. (2013) Short-Read Assembly of Full-Length 16S Amplicons Reveals Bacterial Diversity in Subsurface Sediments. PLoS ONE 8(2):e56018.

53. Caporaso JG et al. (2010) PyNAST: a flexible tool for aligning sequences to a template alignment. Bioinformatics 26(2):266–7

54. Peng Y, Leung HCM, Yiu SM, Chin FYL. (2012) IDBA-UD: a de novo assembler for single-cell and metagenomic sequencing data with highly uneven depth. Bioinformatics 28(11):1420–8.

55. Hyatt D, et al. (2010) Prodigal: prokaryotic gene recognition and translation initiation site identification. BMC Bioinformatics 11:119.

56. Suzek BE, Huang H, McGarvey P, Mazumder R, Wu CH (2007) UniRef: comprehensive and non-redundant UniProt reference clusters. Bioinformatics 23(10):1282–8.

57. Dick GJ, et al. (2009) Community-wide analysis of microbial genome sequence signatures. Genome Biol 10(8):R85.

